# A cross-organism framework for supervised enhancer prediction with epigenetic pattern recognition and targeted validation

**DOI:** 10.1101/385237

**Authors:** Anurag Sethi, Mengting Gu, Emrah Gumusgoz, Landon Chan, Koon-Kiu Yan, Joel Rozowsky, Iros Barozzi, Veena Afzal, Jennifer Akiyama, Ingrid Plajzer-Frick, Chengfei Yan, Catherine Pickle, Momoe Kato, Tyler Garvin, Quan Pham, Anne Harrington, Brandon Mannion, Elizabeth Lee, Yoko Fukuda-Yuzawa, Axel Visel, Diane E. Dickel, Kevin Yip, Richard Sutton, Len A. Pennacchio, Mark Gerstein

**Author notes:** Both authors contributed equally to this work (Anurag Sethi, Mengting Gu).

## Abstract

Enhancers are important noncoding elements, but they have been traditionally hard to characterize experimentally. Only a few mammalian enhancers have been validated, making it difficult to train statistical models for their identification properly. Instead, postulated patterns of genomic features have been used heuristically for identification. The development of massively parallel assays allows for the characterization of large numbers of enhancers for the first time. Here, we developed a framework that uses *Drosophila* STARR-seq data to create shape-matching filters based on enhancer-associated meta-profiles of epigenetic features. We combined these features with supervised machine learning algorithms (e.g., support vector machines) to predict enhancers. We demonstrated that our model could be applied to predict enhancers in mammalian species (i.e., mouse and human). We comprehensively validated the predictions using a combination of *in vivo* and *in vitro* approaches, involving transgenic assays in mouse and transduction-based reporter assays in human cell lines. Overall, the validations involved 153 enhancers in 6 mouse tissues and 4 human cell lines. The results confirmed that our model can accurately predict enhancers in different species without re-parameterization. Finally, we examined the transcription-factor binding patterns at predicted enhancers and promoters in human cell lines. We demonstrated that these patterns enable the construction of a secondary model effectively discriminating between enhancers and promoters.

## Introduction

Enhancers are gene regulatory elements that activate expression of target genes from a distance [1]. Enhancers are turned on in a space- and time-dependent manner, contributing to the formation of a large assortment of cell types with different morphologies and functions, even though each cell in an organism contains a nearly identical genome [2-4]. Moreover, changes in the sequences of regulatory elements are thought to play a significant role in the evolution of species [5-9]. Understanding enhancer function and evolution is currently an area of great interest because many variants within distal regulatory elements also have been associated with various traits and diseases during genome-wide association studies [10-12]. However, the vast majority of enhancers and their spatiotemporal activities remain unknown because of the difficulty in predicting their activity based on DNA sequence or chromatin state [13, 14].

Traditionally, regulatory activity of enhancers and promoters was experimentally validated using low-throughput heterologous reporter constructs, leading to a relatively small number of enhancers that are functionally validated in several selected mammalian cell types [15, 16]. These validated enhancers were typically in conserved noncoding regions [17, 18] with particular patterns of chromatin [19], transcription factor (TF) binding [20], or noncoding transcription [21]. When complex computational methods for predicting tissue/cell line-specific enhancers were trained on these validated enhancers, they could be susceptible to potential biases and were difficult to generalize to other tissues or species as the training data were usually not large enough. Some published methods also trained their model based on TF binding sites [20, 22-24]. The TF binding sites provide a large dataset for training, but introduce obvious bias as most enhancers do not bind to one or a small group of TFs. In addition, it has remained challenging to assess the performance of different methods for enhancer prediction with a limited number of putative enhancers being validated.

Recently, due to the advent of next-generation sequencing, a number of transfection and transduction-based assays were developed to experimentally test the regulatory activity of thousands of regions simultaneously in a massively parallel fashion [25-31]. In these experiments, plasmids each containing a potential enhancer element and a downstream luciferase or green fluorescent protein (GFP) gene are transfected or transduced into cells. The differences in activity of the tested regions are reflected in the differences of the gene expressions as measured by the fluorescence strength. STARR-seq was one such massively parallel reporter assay (MPRA) that was used to test the regulatory activity of the *Drosophila* genome by inserting candidate fragments from the genome within the 3’ untranslated region of the luciferase gene, leading to the identification of thousands of cell type-specific enhancers and promoters [25, 32]. The result of this assay confirmed that active enhancers and promoters tend to be depleted of histone proteins and contain accessible DNA where various TFs and cofactors bind [33, 34]. In addition, it showed that the regulatory regions tend to be flanked by nucleosomes that contain histone proteins with certain characteristic post-translational modifications.

These attributes lead to an enriched peak-trough-peak (“double peak”) signal in different ChIP-Seq experiments for various histone modifications such as acetylation on H3K27 and methylations on H3K4. The troughs in the double peak ChIP-seq signal represent the accessible DNA that leads to a peak in the DNase-I hypersensitivity (DHS) at the enhancers [35]. These patterns revealed from aggregation of STARR-seq-validated enhancers provide a new scope for annotating regulatory regions. The large amount of experimental data also allows us to train complex models with less bias and test the performance of different models using cross validation.

Here we developed a framework for making supervised enhancer prediction models using MPRA datasets. We made use of published data resources to provide a comprehensive model for enhancer prediction that can be applied across different contexts (i.e., different species and tissue types), and we validated our model in a variety of contexts. In particular, we utilized extensive datasets from STARR-seq experiments performed on *Drosophila* cell lines to create and parameterize our model. Unlike previous prediction methods that focused on the enrichment (or signal peaks) of different epigenetic datasets, we developed a method to take into account the specific enhancer-associated pattern within different epigenetic signals. As the epigenetic signal around each enhancer is noisy, we aggregated the signal around thousands of enhancers identified using STARR-seq to increase the signal-to-noise ratio and identify the shape associated with active regulatory regions. Previous ENCODE and modENCODE efforts showed that the chromatin modifications on active promoters and enhancers are conserved across higher eukaryotes [36-42]. The signals of different chromatin modifications upstream of genes were used to create a universal model for predicting gene expressions; moreover, the parameters of the model were transferable across humans, flies, and worms. Here, we further explored this conservation of epigenetic signal shapes for constructing simple-to-use transferrable statistical models with six parameters to predict enhancers and promoters in diverse eukaryotic species including fly, mouse, and human. We showed that the enhancer predictions from our transferrable model are comparable to the prediction accuracy of species-specific models.

Working across organisms also allowed us to take advantage of different assays to validate our predictions in a robust fashion using multiple experimental approaches. In the first stage, we predicted enhancers in six different embryonic mouse tissues and tested the activity of these predictions *in vivo* with transgenic mouse assays. Due to the ethical considerations of performing such transgenic assays in human embryos, we then proceeded to test the activity of these elements in a human cell line *in vitro,* e.g. H1 human embryonic stem cells (H1-hESCs), an extensively studied and well-characterized cell line.

After validating our predictions, we examined the TF binding patterns in enhancers and promoters respectively. The comprehensive set of TF binding experiments available for H1-hESC cells enabled us to perform this analysis. We found that the pattern of TF and co-TF binding at active enhancers is much more heterogeneous than the corresponding patterns on promoters, which can be used to distinguish enhancers from promoters with high accuracy. Thus, our methods provide a framework that utilizes different whole-genome epigenetic datasets to predict active regulatory regions in a cell type-specific manner. Further functional genomics datasets could be utilized to identify key TFs associated with active regulatory regions within these cell types.

## Results

### Aggregation of epigenetic signal in *Drosophila* to create a metaprofile

We developed a framework to predict active regulatory elements using the epigenetic signal patterns associated with experimentally validated promoters and enhancers. We aggregated the signal of histone modifications on STARR-seq peaks to remove noise in the signal and created a metaprofile of the double peak pattern of histone modifications flanking enhancers and promoters. STARR-seq peaks typically consist of a mixture of enhancers and promoters, and at this stage we did not differentiate between the two sets of regulatory elements. As STARR-seq quantifies enhancer activity in an episomal fashion, not all peaks would be active in the native chromatin environment. Arnold and colleagues showed that the STARR-seq peaks that occur with enriched DNase hypersensitivity or H3K27ac modifications tend to be associated with active genes, whereas other STARR-seq peaks tend to be associated with enrichment of repressive marks such as H3K27me3 [25]. Hence, we took the overlap of the STARR-seq enhancers with H3K27ac and/or DHS peaks to get a high confidence set of enhancers that are active *in vivo*, based on which the metaprofiles were created. These metaprofiles were then utilized in a pattern recognition algorithm for predicting active regulatory elements in a cell type-specific manner.

The STARR-seq studies on *Drosophila* cell lines provide the most comprehensive datasets as they were performed genome-wide and performed with multiple core promoters [25, 43]. We created metaprofiles using the histone modifications and DNase signals at active STARR-seq peaks (see Fig. 1 and Methods) identified within the *Drosophila* S2 cell line. Approximately 70% of the active STARR-seq peaks contained an easily identifiable double peak pattern even though there was variability in the distance between the two maxima of the double peak in the ChIP-chip signal (Supplementary Fig. 1). While the minimum tended to occur in the center of these two maxima on average, the distance between the two maxima in the double peaks varied between 300 and 1,100 base pairs. During aggregation, we first aligned the two maxima in the H3K27ac signal across different STARR-seq peaks, followed by interpolation of the signal before calculating the average metaprofile. We performed a flipping step to generate the matched filter for convolution, which maintains the asymmetry in the underlying H3K27ac double peak because it may be associated with the directionality of transcription [44]. Then we calculated the dependent metaprofiles for 30 other histone marks by applying the same set of transformations to these datasets. These dependent metaprofiles also exhibited a double peak pattern, and the maxima across different histone modification signals tended to align with each other on average (Supplementary Fig. 2). This indicates that a large number of histone modifications tends to simultaneously co-occur on the nucleosomes flanking an active enhancer or promoter. In contrast, as expected, the DHS signal displayed a single peak at the center of the H3K27ac double peak (Fig. 1). In addition, repressive marks such as H3K27me3 were depleted in these regions, and the metaprofile for these regions did not contain a double peak signal (Supplementary Fig. 2).

**Figure 1:**
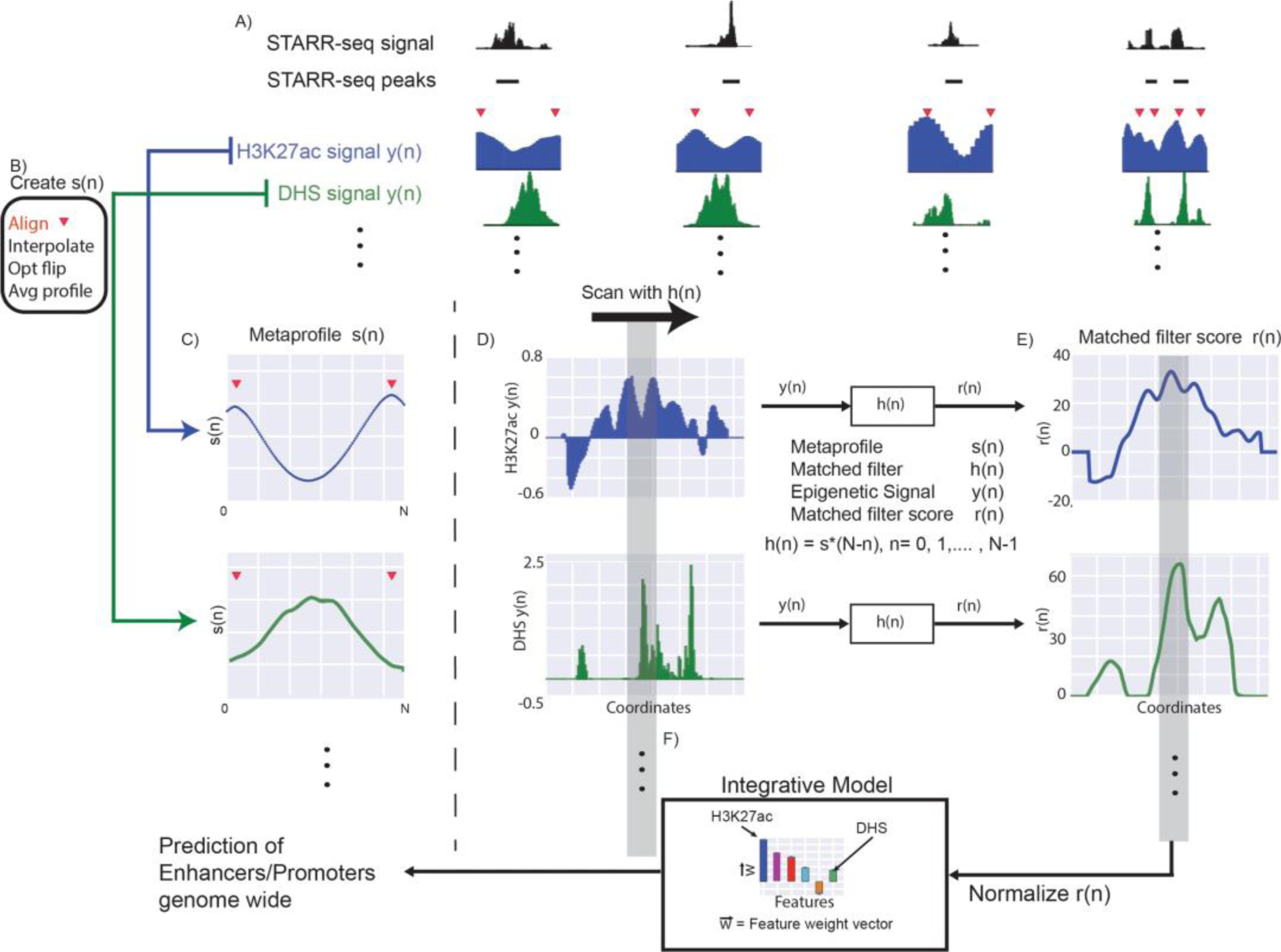
Flowchart of the Matched-filter model. A) We identified the “double peak” pattern in the H3K27ac signal close to STARR-seq peaks. The red triangles denote the position of the two maxima in the double peak. B) We aggregated the H3K27ac signal around these regions after aligning the flanking maxima, using interpolation and smoothing on the H3K27ac signal, and averaged the signal across different MPRA peaks to create the metaprofile in C). The same operations were performed on other histone signals and DHS to create metaprofiles in other dependent epigenetic signals. D) Matched filters were used to scan the histone and/or DHS datasets to identify the occurrence of the corresponding pattern in the genome. E) The matched filter scores are high in regions where the profile occurs (grey region shows an example) but low when only noise is present in the data. The individual matched filter scores from different epigenetic datasets were combined using integrated model in F) to predict active promoters and enhancers in a genome-wide fashion.

### Match of a metaprofile is predictive of regulatory activity

We evaluated whether these metaprofiles could be utilized to predict active promoters and enhancers using matched filters, a well-established algorithm in template recognition. Matched filter is the pattern recognition algorithm that uses a shape-matching filter to recognize the occurrence of a template in the presence of stochastic noise [45]. When scanning the whole genome with matched filter, we applied the H3K27ac metaprofile to match the region between histone double peaks. Due to the aforementioned variability of the distance between the double peaks, we allowed the width of the scanned regions to vary between 300 and 1,100 basepairs. We used the highest score to rate the regulatory potential of this region (see Methods). The dependent profiles were subsequently used on the region of the same width to get the score for other histone marks.

We calculated the matched filter score for all 30 epigenetic modification signals available in the *Drosophila* cell lines on STARR-seq peaks and a negative control set (Supplementary Fig. 3). Interestingly, the distribution of matched filter scores for STARR-seq peaks are unimodal for each histone mark except for H3K4me1, H3K4me3, and H2Av, which are bimodal. We looked at the degree to which the matched filter scores for promoters and enhancers are higher than the matched filter scores for the rest of the genome, as this is a measure of the signal-to-noise ratio for regulatory region prediction. We found that, in general, several histone acetylations marks including H3K27ac, as well as H1, H3K4me2, and DHS were the most accurate prediction features, whereas other histone marks like H3K79m1 and H4K20me1 were not well suited as their matched filter scores for positive regions and negative regions were not distinguishable.

To quantitatively evaluate whether the occurrence of the epigenetic metaprofiles could be used to predict active enhancers and promoters, we did a ten-fold cross validation assessing the average areas under the receiver operating characteristic (ROC) (AUROC) and the areas under the precision-recall (PR) (AUPR) curves. PR curves are particularly useful to assess the performance of classifiers in skewed or imbalanced data sets in which one of the classes is observed much more frequently compared to the other class. Comparing the matched filter result with the peak calling result, we found that the AUROC and AUPR of the matched filter scores for different histone modifications were higher than those of the peaks of corresponding histone marks (Supplementary Fig. 4), suggesting that the matched filter score is more accurate for predicting active STARR-seq peaks than the simple enrichment of the signals.

We observed that the H3K27ac matched filter score is the most accurate feature for predicting active regulatory regions identified using STARR-seq (Fig. 2 and Supplementary Table 1). This could potentially be a bias as we used STARR-seq peaks overlapping with H3K27ac or DHS peaks as the positive training set as described above. To check for this, we repeated the analysis using all STARR-seq peaks as the positives. We found that H3K27ac matched filter scores still had the highest AUROC and AUPR compared to the other features (Supplementary Fig. 5). In addition, while DHS peaks were used to select the STARR-seq peaks, the AUROC and AUPR for DHS did not seem to be as high as some other histone marks. Thus, the selection of STARR-seq peaks using H3K27ac and DHS was less likely to have introduced substantial bias to the model, but they helped to prune the STARR-seq peaks to create a more accurate set for training.

**Figure 2:**
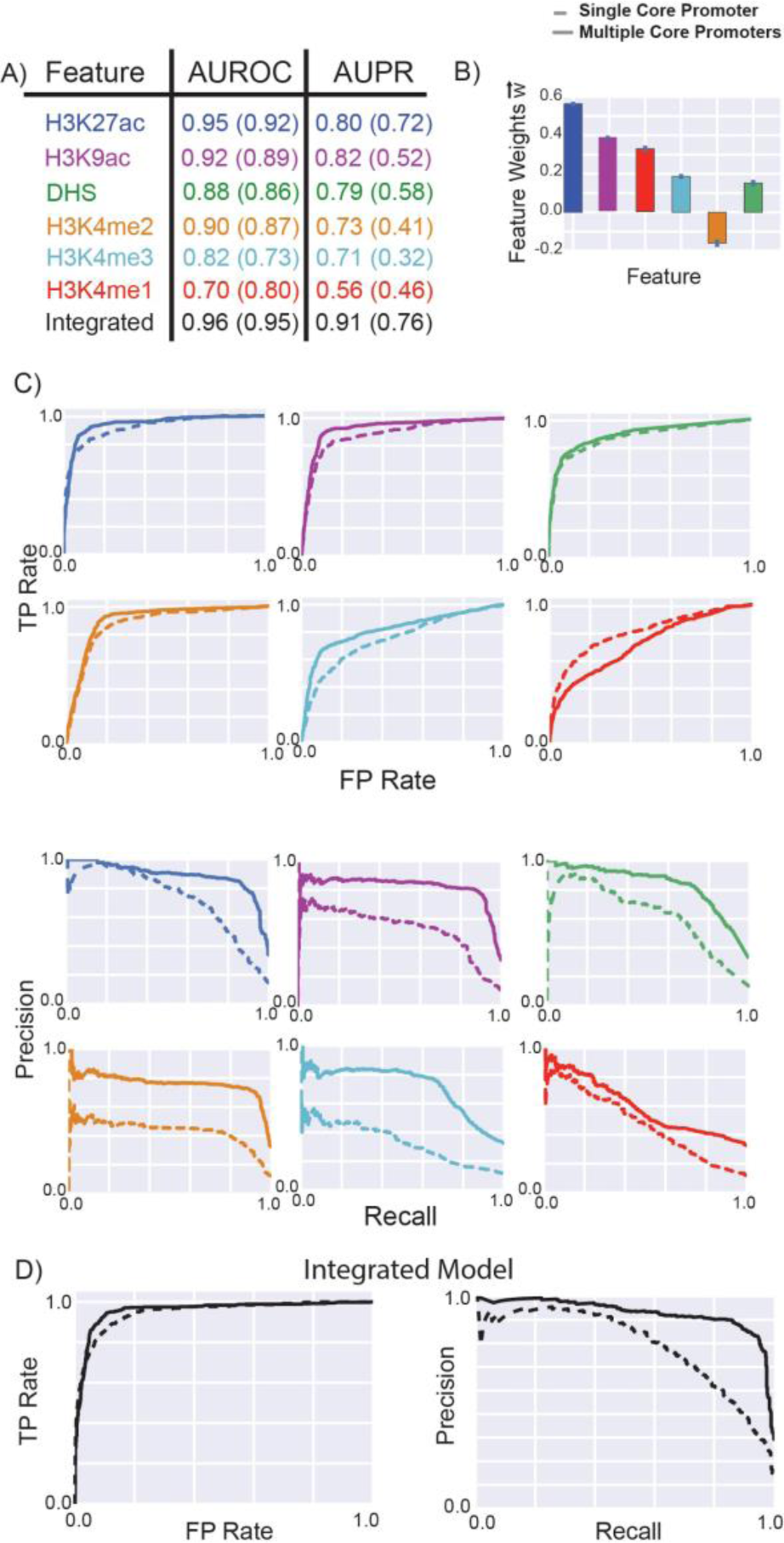
Performance of matched filters and integrated models for predicting MPRA peaks. The performance of the matched filters of different epigenetic marks and the integrated model for predicting all STARR-seq peaks was compared using ten-fold cross validation. A) The area under the receiver-operating characteristic (AUROC) and the precision-recall (AUPR) curves were used to measure the accuracy of different matched filters and the integrated model. B) Weights of the different features in the integrated model are plotted; error bars show the standard deviation of feature weights measured by ten-fold cross validation. These weights may be used as a proxy for the importance of each feature in the integrated model. C-D) The individual ROC and PR curves for each matched filter and the integrated model are shown. The performance of these features and the integrated model for predicting the STARR-seq peaks using multiple core promoters and a single core promoter were compared. The numbers within the parentheses in A) refer to the AUROC and AUPR for predicting the peaks using a single STARR-seq core promoter; the numbers outside the parentheses refer to the performance of the model for predicting peaks from multiple core promoters.

As STARR-seq identified active regulatory regions display core-promoter specificity --different sets of enhancers are identified when different core promoters are used in the same cell type [43, 46-49], we combined the peaks identified from multiple STARR-seq experiments of S2 cells and reassessed the performance of the matched filters at predicting these regulatory regions. Merging the STARR-seq peaks from multiple core promoters used in S2 cells led to higher AUROC and AUPR for the matched filters of most histone marks (Fig. 2).

### Machine learning can combine matched filter scores from different epigenetic features

We built an integrated model with combined matched filter scores of six commonly available and discriminative epigenetic marks (H3K27ac, H3K4me1, H3K4me2, H3K4me3, H3K9ac, and DHS) associated with active regulatory regions using a linear support vector machine (SVM) [50]. The selection of these six features was based on their matched filter score performance, their importance in the integrated model, and data availability. The performance of each individual feature was measured by how well its matched filter score could distinguish STARR-seq peaks from negative controls, as discussed above (Supplementary Fig. 3 and Supplementary Table 1). The importance of each feature in the integrated model was measured by the feature coefficient or GINI score.

We combined the matched filter scores from all 30 measured histone marks along with the DHS in statistical models like random forest and SVM (Supplementary Fig. 6). The integrated models with 30 epigenetic features displayed high accuracy (AUROC=0.97, AUPR=0.93 for SVM model with multiple core promoters). We found that several major histone modification marks had considerably high weights in these integrative models, like histone acetylations (H3K27ac, H3K9ac, H2BK5ac, H4ac, and H4K12ac) and H3K4 methylations (H3K4me1, H3K4me2, H3K4me3). While the DHS performed well as on its own in the single-feature matched filter model (Fig. 2), it had a lower weight in the integrated SVM among the six features, likely due to the fact that the information in DHS is redundant with the information contained within the histone mark (e.g., the DHS peaks usually occur at the trough region between two maxima in the histone signal). Despite the redundancy, the combination of the DHS and histone signals was more predictive of regulatory activity because the reinforcing signals strengthened the prediction as compared to the uncorrelated noise.

We then checked the data availability of the features selected from above. As our goal was to build a model with broad applicability across organisms, we excluded epigenetic marks that were generally unavailable in mammalian tissues and cell lines (e.g., H2BK5ac, H4ac and H4K12ac). The final model consisted of a very simplified set of input: H3K27ac, H3K4me1, H3K4me2, H3K4me3, H3K9ac, and DHS. We then assessed the performances of different statistical approaches including random forest, ridge regression and Naïve Bayes and SVM to combine the features. While all these approaches performed similarly (Supplementary Fig. 7), we used a linear SVM in our framework due to its better interpretability.

We found that the simplified SVM model had a high performance similar to that of the complete SVM model using all 30 epigenetic marks, with an AUROC of 0.96 (0.97 in the full model) and an AUPR of 0.91 (0.93 in the full model). We observed that H3K27ac was the highest weighted feature in SVM. To test if this was biased because of the selection of STARR-seq peaks based on H3K27ac and DHS, we trained an SVM model using all STARR-seq peaks with the same six features. We found that H3K27ac still had the highest GINI score in random forest, albeit a slightly smaller coefficient in SVM (Supplementary Fig. 8), which could be due to the added noise in the training data. In general, the integrated model trained on the six features achieved good performance upon cross-validation, and this set of input features allowed the integrated model to be applied to a variety of cell lines and tissues, as many relevant ChIP-seq and DNase experiments have been performed by the Roadmap Epigenomics Mapping [51] and the ENCODE [52] Consortia in a wide variety of samples.

We repeated this analysis training a six-parameter integrative models using STARR-seq data in the *Drosophila* BG3 cell line, and tested them with STARR-seq data in the S2 cell line. Again, we found that these statistical models showed similar high prediction accuracy in predicting enhancers and promoters in the S2 cell line (Supplementary Fig. 9). This result indicates that our framework of combining epigenetic features with a linear SVM model to predict enhancers is applicable across different cell lines.

To evaluate the impact of the training sample size on model performance, we did a saturation analysis where we down-sampled the training data to different levels of fractions and evaluated the model performance on the remaining data. For each fraction level, we did a ten-fold cross-validation (see methods) and then took the average of the ten outputs. We found that the average AUPR increased with increasing size of training data, and started to saturate for our SVM model with 80-90% of the experimental data for training (Supplementary Fig. 10). The average AUROC remained comparable, although the variances decreased with increasing training data size. This indicates that a five-fold cross validation might be sufficient with this size of data, as a five-fold cross validation uses 80% of the data for training and the remaining 20% of the data for testing, at which point the performance of the model started to saturate.

### Distinct epigenetic signals associated with promoters and enhancers

We proceeded to create individual metaprofiles and machine learning models for the two classes of regulatory activators – promoters (or proximal) and enhancers (or distal). While strictly discriminating enhancers from promoters is very difficult, one commonly used strategy in the published literature is to look at their distance to the closest transcription start site (TSS). In conjunction to that, we divided all the active STARR-seq peaks into promoters or enhancers based on this distance to delineate their likely function in the native context. Due to the conservative distance metric used in this study (1kb upstream and downstream of TSS in *Drosophila* genome), the enhancers are regulatory elements that are not close to any known TSS and could be considered to enhance gene transcription from a distance. However, a few of the promoters may also regulate distal genes in addition to their promoter activity. We then created metaprofiles of the different epigenetic marks on the promoters and enhancers and assessed the performance of the matched filters for predicting active regulatory regions within each category (Fig. 3). We also combined the peaks identified from multiple STARR-seq experiments of S2 cells and reassessed the performance of the matched filters at predicting promoters and enhancers, respectively. Merging the STARR-seq peaks from multiple core promoters led to higher AUROC and AUPR for the matched filters of most histone marks (Supplementary Table 2). The highest matched filter scores were typically observed on promoters, and the matched filters for each of the six features tended to perform better for promoter prediction. Similar to previous studies [53, 54], we observed that the H3K4me1 metaprofile was very predictive for enhancers but was close to random for predicting promoters. In contrast, the H3K4me3 metaprofile could be utilized to predict promoters and not enhancers. The histogram for matched filter scores showed that the H3K4me1 matched filter score was higher near enhancers while the H3K4me3 matched filter score tended to be higher near promoters (Supplementary Fig. 11). The mixture of these two populations led to bimodal distributions for H3K4me1 and H3K4me3 matched filter scores when calculated over all regulatory regions (Supplementary Fig. 3).

**Figure 3:**
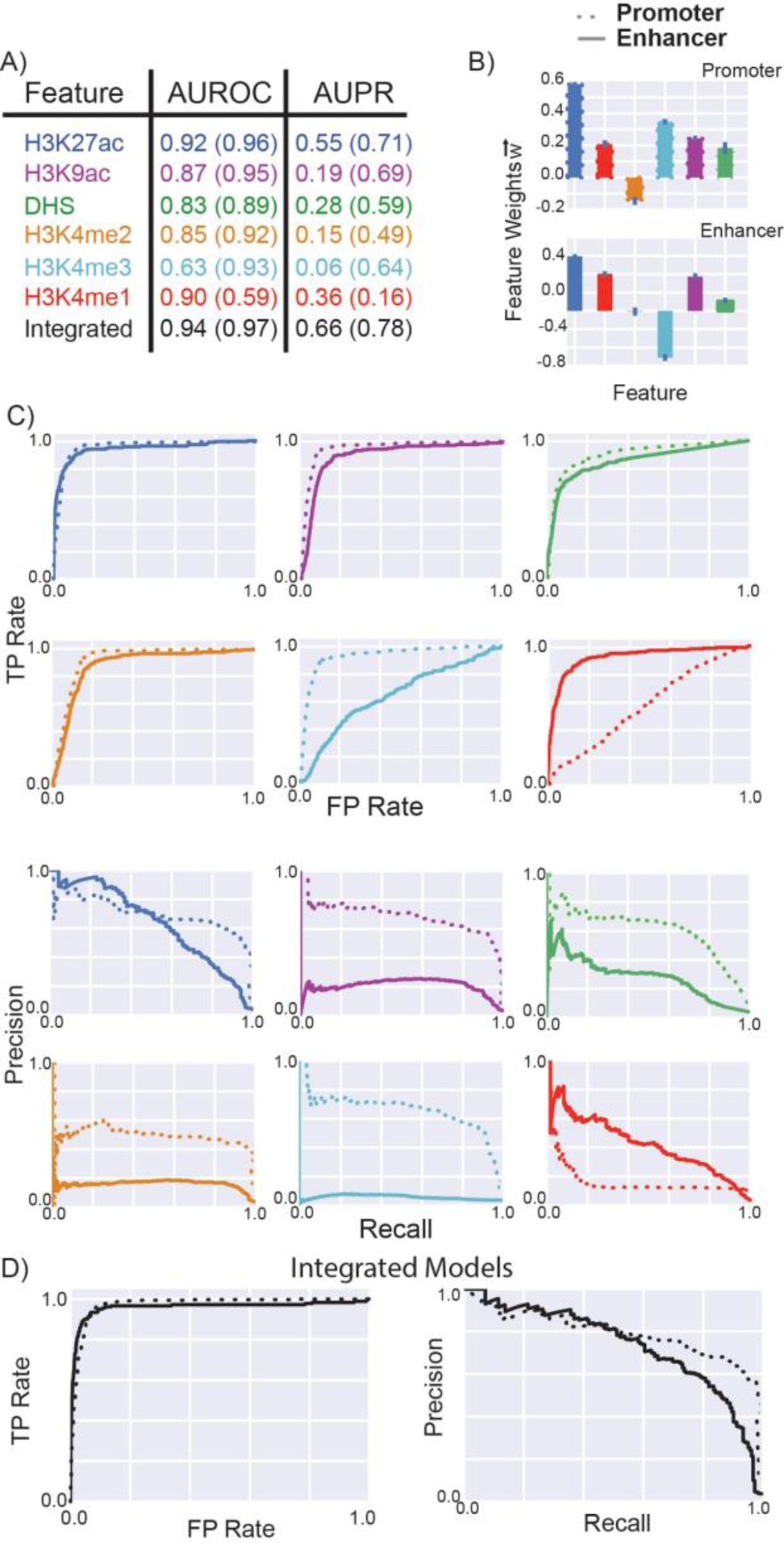
Performance of matched filters and integrated models for predicting promoters and enhancers. The performance of the matched filters of different epigenetic marks and the integrated model for predicting active promoters and enhancers were compared using ten-fold cross validation. A) The numbers within parentheses refer to the AUROC and AUPR for predicting promoters; the numbers outside the parentheses refer the performance of the models for predicting enhancers. B) Weights of the different features in the integrated models for promoter and enhancer prediction are plotted; error bars show the standard deviation of feature weights measured by ten-fold cross validation. C-D) The ROC and PR curves for each matched filter and the integrated model are shown. The performance of these features and the integrated model for predicting the active promoters and enhancers using multiple core promoters were compared.

We again trained different statistical models to learn the combination of features associated with promoters and enhancers respectively. These integrated models outperformed the individual matched filters at predicting active enhancers and promoters (Fig. 3 and Supplementary Fig. 12). In addition, the weights of the individual features identified the difference in the roles of H3K4me1 and H3K4me3 matched filter scores at discriminating active promoters and enhancers from inactive regions in the genome. The promoter-based (enhancer-based) model performed much more poorly at predicting enhancers (promoters) indicating the unique properties of these regions (Supplementary Figs. 13 and 14). We also created two integrated models utilizing matched filter scores of all 30 histone marks as features for predicting enhancers and promoters. The additional histone marks provided independent information regarding the activity of promoters and enhancers as these features increased the accuracy of these models (Supplementary Fig. 15). The weights of different features indicated that H2BK5ac again displayed the most independent information for accurately predicting active enhancers and promoters. We observed similar trends and accuracy with several different machine learning methods (Supplementary Figs. 6 and 15). To investigate the impact of different distance metrics used to segregate enhancers and promoters, we repeated our analysis with different distance cutoffs (0.5kb, 1.5kb, 2.0kb and 2.5kb). While the accuracy as measured by the AUROC of different features and the integrated model slightly reduced as the distance increased, the importance of each feature in the integrated model remained similar (Supplementary Fig. 16).

### Application of the STARR-seq model to predict enhancers in mammalian species

One of the important findings of previous ENCODE and model organism ENCODE efforts was the conservation of chromatin marks close to regulatory elements across hundreds of millions of years of evolution [36-42]. The relationship of chromatin marks to gene expression was very similar, for instance, in worms, flies, mice and humans, so much that one could build a statistical model relating chromatin modification to gene expression that would work without re-parameterization across different organisms. This motivated us to apply our well-parameterized model based on the STARR-seq data from flies to mammalian systems, eg. mouse and human, and test our model performance.

We started by making genome-wide predictions of regulatory regions in mouse. We processed tissue-specific epigenetic signals and used them in our model to account for the tissue specificity of enhancers. Predictions were made in six different tissues (forebrain, midbrain, hindbrain, limb, heart and neural tube) at the embryonic day 11.5 (e11.5) stage (Genome-wide predictions are available through our website at https://goo.gl/E8fLNN). These tissues were selected as their epigenetic signals have been highly studied in mouse ENCODE, providing us with a rich source of raw data that could be utilized for making enhancer and promoter predictions. In addition, the VISTA database contains close to 100 validated enhancers that could be used to test predictions in each of these tissues. Using our model, we predicted 31K to 39K regulatory regions in individual tissues in mouse, with each region ranging from 300bp to 1,100bp. Notably, a consistent proportion of two-thirds (66-70%) of these predicted regulatory regions were distal regulatory elements for all six tissues, with the other one-third (30-34%) being proximal regulators (Supplementary Table 3). These numbers agree with a previous enhancer evolution study [8], and suggest that the amount of enhancers and promoters are likely comparable in different tissues.

Similarly, we performed a genome-wide prediction of regulatory regions in the ENCODE top-tier human cell lines, including H1-hESC, GM12878, K562, HepG2, A549 and MCF- 7 (all available through our website). For each cell line, we utilized the six-parameter integrated model to predict active enhancers and promoters based on the epigenetic datasets measured by the ENCODE consortium [52]. In H1-hESC, for example, we predicted 43,463 active regulatory regions, of which 22,828 (52.5%) were within 2kb of the TSS and were labeled as promoters. A large proportion of the predicted enhancers were found in the introns (30.41%) and intergenic regions (13.93%) (Supplementary Fig. 17). The predicted promoters and enhancers were significantly closer to active genes than expected randomly (Supplementary Fig. 18).

### Whole-genome STARR-seq enables proper training of enhancer prediction model

We next evaluated how well the STARR-seq model could predict mammalian enhancers. Particularly, we wanted to compare the current mouse enhancer predictions with predictions from models directly trained on mouse data. The relatively large number of known mouse enhancers from the VISTA database enabled us to parameterize a model in the same way as we did with the *Drosophila* STARR-seq data. However, the number of active enhancers in each tissue in the VISTA database is not at the same scale as the STARR-seq dataset. In total, we found 1,253 positive regions and 8,631 negative regions pulling together from different tissues.

With the VISTA database, we trained four models based on four sets of available e11.5 mouse tissue-specific enhancers (hindbrain, limb, midbrain and neural tube), and assessed them using ten-fold cross-validation. There were no DHS data available for e11.5 forebrain and heart, thus these two tissues were excluded for fair comparison. The average AUROC value was compared to the AUROC of testing the STARR-seq trained model on the same VISTA enhancer data. Despite the significantly unbalanced negative-to-positive ratios of mouse enhancers in the database, the six-parameter integrative SVM models learned using balanced *Drosophila* STARR-seq data were highly accurate at predicting active enhancers and promoters in mouse (Supplementary Fig. 19a). The cross-validated mouse model, although it did well, performed no better on predicting mouse tissue-specific enhancers. We found that the best performing one among the mouse models was for the tissue midbrain, likely due to the fact that the number of validated midbrain enhancers is the largest. To construct a larger training sample for mouse, we pooled together the normalized z-scores of matched filter scores for six epigenetic signals of all four tissues, and parameterized a model using this larger set of data. Again, we observed that the original model trained with *Drosophila* STARR-seq data performed equally well at predicting mouse enhancers and much better in predicting fly enhancers (Supplementary Fig. 19b). Overall, the result suggests that using the larger and more comprehensive STARR-seq data set for parameter tuning is superior to using the smaller mouse data set, even on mouse.

Given the above overall statistical evaluations, we are confident in the STARR-seq parameterized model. We then set out to do targeted unbiased validations of the mammalian enhancers predicted, which is described in the next two sections.

### Validation *in vivo* in mouse

To test the activity of predicted mouse enhancers *in vivo*, we performed transgenic mouse enhancer assays in e11.5 mice for 133 regions, including 102 regions selected based on the H3K27ac signals rank of corresponding mouse tissues, and 31 regions selected by an ensemble approach from human homolog sequences. For each tested candidate, a readout of activity across the entire embryo was collected. The number of transgenic mice that showed the pattern for each tissue was also recorded to confirm reproducibility (See Methods, Supplementary Table 4 and 5, and Supplementary Fig. 20). In addition, we included other published transgenic mouse experiments from the VISTA database for validation. This large set of validated enhancers allowed us to comprehensively evaluate our enhancer predictions in all six e11.5 mouse tissues.

Among the first 102 tested regions, 62 were selected based on forebrain H3K27ac signal rank, with 20, 22, and 20 regions being in the top, middle and bottom rank, respectively. We found that while the top-ranked regions showed high activity rate (~70%), the active rate for the middle rank and the bottom rank is similar, with a slightly higher active rate for the latter (32% and 35%). This might suggest that the simple H3K27ac signal rank is not a direct indicator of enhancer activity. Another 40 regions were selected by heart H3K27ac signal with half of them coming from the top rank and the other half coming from the middle rank. We observed 35% of the top rank regions and 25% of the middle rank regions showing enhancer activity. For the other 31 human homolog sequences, 12.9% and 9.7% of the assessed regions were active in heart and forebrain, respectively. The lower active rate was likely due to the fact that these human sequences are less well behaved in mouse tissues compared to their original native environment.

We evaluated the predictability of our matched filter model for each individual histone marks and DHS, as well as the integrated SVM model (Fig. 4). For each tissue, our model ranked all the tested candidate elements with their predicted activity in this tissue using either individual features or the integrated SVM model. Then, the label of each element from experiment readout was used to assess the predictions with ROC and PR curves. On average, the integrated model trained with *Drosophila* STARR-seq data achieved an AUROC of 0.80 and an AUPR of 0.37 for tissue-specific enhancer predictions in mouse (Fig. 4a). For AUROC, the baseline was always 0.50, whereas for AUPR the baseline was the positive rate from the experiment. In our transgenic mouse experiment, the positive rate varied from 8.8% to 17.6% among the tissues, and thus the AUPR had a larger variance compared to the AUROC. We also did a similar evaluation using the regulatory elements identified by the transduction-based FIREWACh assay in mouse embryonic stem cells (mESCs) [30]. Again, we observed similar results for individual histone marks and combined SVM model (Supplementary Fig. 21). As the *in vivo* and FIREWACh assays utilized a single core promoter to validate regulatory regions, the performance of the different models in Figures 4 and Supplementary Figure S21 are probably underestimated.

**Figure 4:**
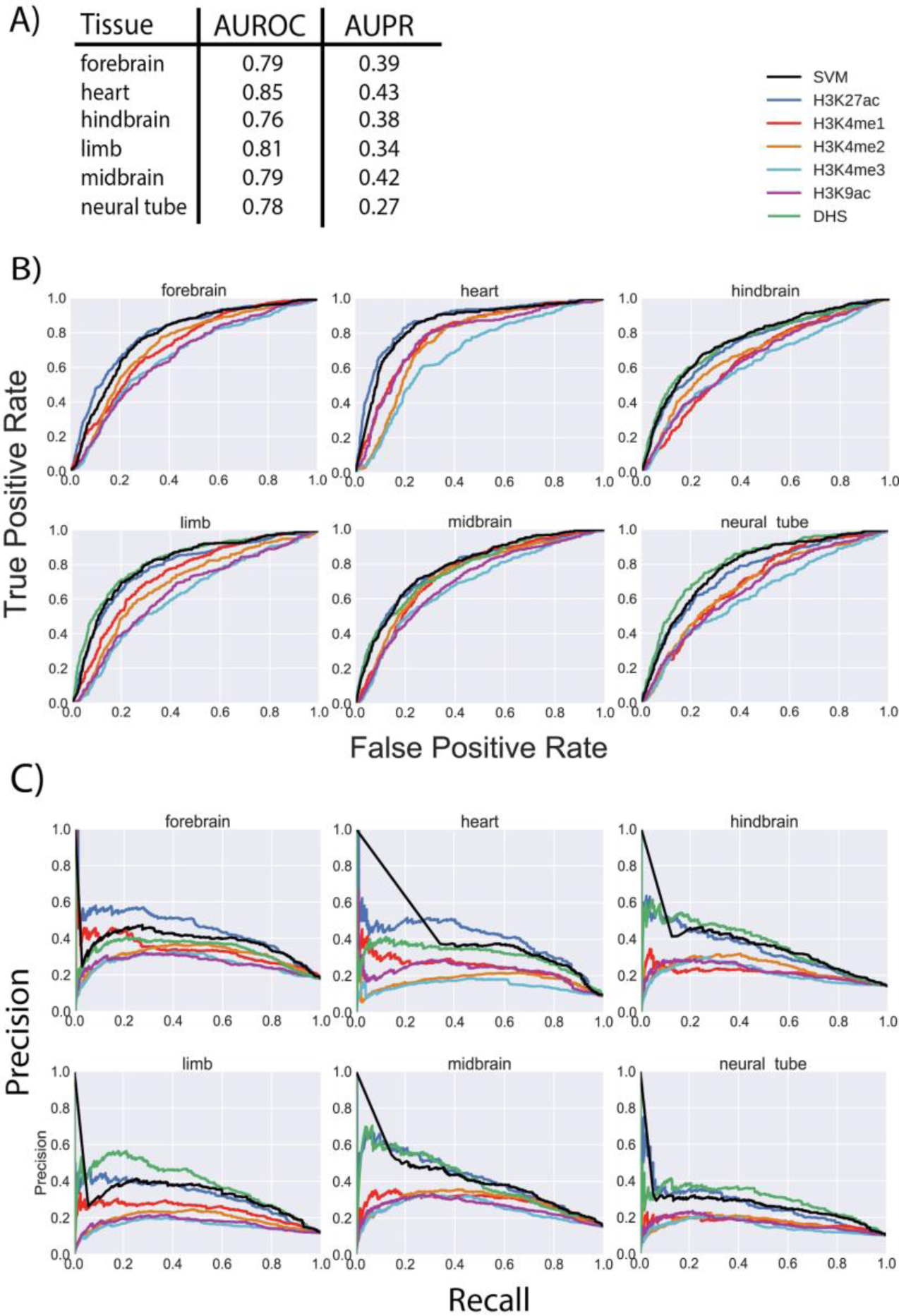
Performance of matched filters and integrated model for predicting active enhancers in mice. The performance of the *Drosophila* STARR-seq-based matched filters and the integrated model for predicting active enhancers identified by transgenic mouse enhancer assays in six different tissues of e11.5 mice. A) The AUROC and AUPR are shown for the integrated SVM model in six tissues. The weights of the different features in the integrated model are the same as the weights shown in Figure 3 for enhancers. B) The individual ROC curves of each feature and the integrated SVM model for each tissue are shown. C) The individual PR curves of each feature and the integrated SVM model for each tissue are shown.

### Validation in human cell lines

We proceeded to validate our STARR-seq based model for predicting human enhancers using a cell-based transduction assay. We used a third-generation, self-inactivating (SIN) HIV-1 based vector system in which the enhanced GFP (eGFP) reporter was driven by the DNA element of interest to test putative enhancers after stable transduction of four cell lines, including H1-hESCs (Fig. 5). The predicted enhancers, ranging from 650 to 2,500 bp, were PCR amplified from human genomic DNA and separately inserted immediately upstream of a basal Oct-4 promoter of 142 bp within the SIN HIV vector. Each putative enhancer was tested in triplicate for both forward and reverse orientation in H1-hESCs. We used empty SIN HIV vector and FG12 as the negative and the positive controls, respectively. Note that the empty vector had the basal Oct-4 promoter, along with the IRES-eGFP reporter cassette. We assessed putative enhancer activity by flow cytometric readout of eGFP expression 48-72 h post-transduction, normalized to the negative control.

**Figure 5:**
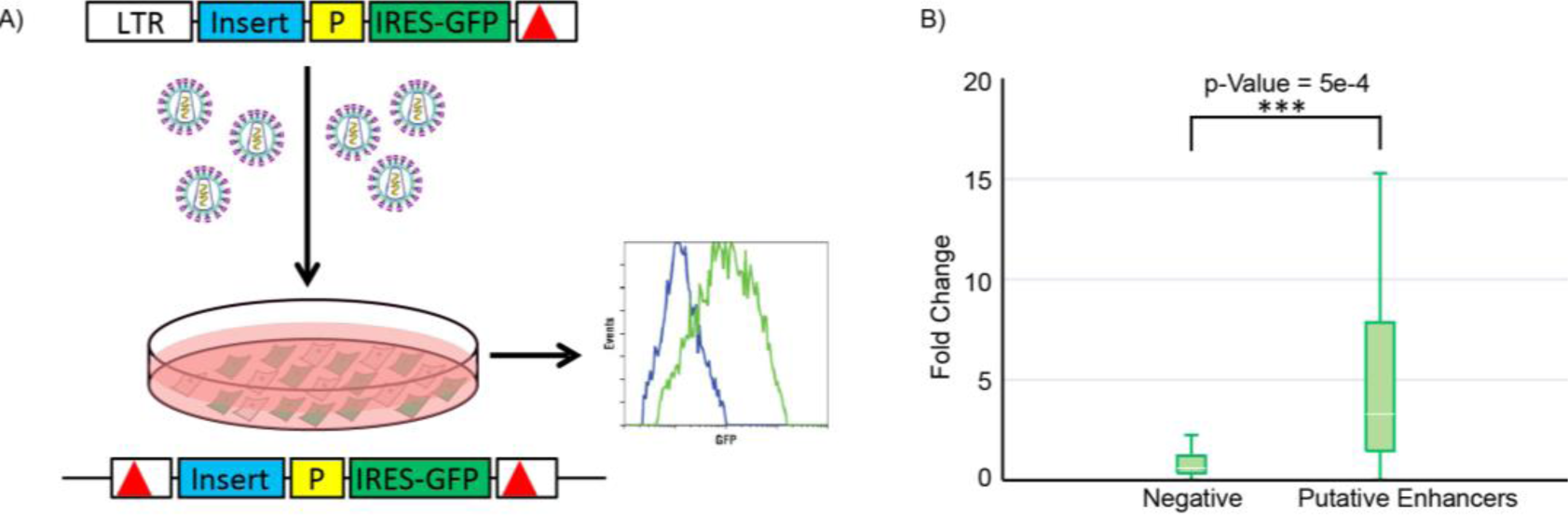
Enhancer validation experiments. A) Schematic of the enhancer validation experiment flow. At top is the third-generation HIV-based self-inactivating vector (deletion in 3’ LTR indicated by red triangle), with PCR-amplified test DNA (blue, cloned in both orientations) inserted just 5’ of a basal Oct4 promoter (P) driving IRES-eGFP (green). Vector supernatant was prepared by plasmid co-transfection of 293T cells. Cells of interest were transduced and then analyzed by flow cytometry a few days later. Shown below is the expected post-transduction structure of the SIN HIV vector, with a duplication of the 3’ LTR deletion rendering both LTRs non-functional. B) Fold changes of gene expression of eGFP was compared between negative elements and putative enhancers chosen at random, with the p-value measured by the Wilcoxon signed-rank test. The 25th and 75th percentiles of the fold change in gene expression for each group are represented by the whiskers in the box plot.

We selected a total of 23 predicted intergenic enhancers for validation. These predictions were chosen at random to ensure that they truly represented the whole spectrum of predicted enhancers and not just the top tier of predicted enhancers. Of these 23 putative enhancers, 20 were successfully PCR-amplified and cloned into the SIN HIV vector in both directions. To measure the distribution of gene expression in the absence of enhancer, we also amplified and cloned 20 non-repetitive elements with a similar length distribution that were predicted to be inactive into the same SIN HIV vector. All positive and negative DNA elements were transduced and tested for activity in both forward and reverse orientations as enhancers are thought to function in an orientation-independent manner. Following the same procedures, we performed functional testing in duplicate in HOS, TZMBL, and A549 cell lines in addition to H1-hESCs.

Insertion of 12 of the putative enhancers into the HIV vector resulted in a significant increase in eGFP expression (P-value < 0.05 over the distribution of gene expression for negative elements) in H1-hESCs (Supplementary Table 6). Most of the positive enhancers displayed a significant increase in gene expression irrespective of their orientation. In contrast, the negatives displayed much lower levels of gene expression (Fig. 5 and Supplementary Fig. 22). The activity of these tested enhancers also showed cell specificity. For example, predicted H1-hESC enhancers A1 to A6 all showed strong activity in both orientations in H1-hESCs (Supplementary Fig. 23), but not in A549, HOS or TZMBL cell lines (Supplementary Fig. 24). Overall, 16 of the 20 tested predictions displayed a statistically significant increase in gene expression of the reporter gene in at least one of the cell lines. Given the promoter specificity of enhancers in such assays, we anticipate that some of the elements that could not be validated in this particular vector would function as enhancers in a more natural biological context (e.g., with the cognate promoter or in the absence of surrounding HIV vector sequences).

### Comparison against other computational methods

To further assess the performance of our model, we compared it against other published methods. We first did a comparison with ChromHMM [55], a well-known method to segment the genome based on chromatin features. We evaluated the performance of ChromHMM in the same way as we presented above using the same data from the transgenic mouse enhancer assay. Our integrated model outperformed ChromHMM in all four tissues, with an AUROC value of 0.76 in hindbrain (versus ChromHMM 0.69), and 0.81 in limb (versus ChromHMM 0.75), etc (Supplementary Fig. 25). In addition to the comparison with unsupervised segmentation-based methods, we also compared with other published enhancer prediction tools, including CSIANN, a neural network based approach [56]; DELTA, an ensemble model integrating different histone modifications [57]; RFECS, a random forest model based on histone modifications [53], and REPTILE, a more recent published method that integrates histone modifications and whole-genome bisulfite sequencing data [58]. We used the mouse experimental data published in REPTILE for the comparison, and assessed the performance of our method compared to the four published methods mentioned above for all four mouse tissues with available experimental data, ChIP-seq data, and DNase data. In three out of four tissues (hindbrain, limb and neural tube), our method had the highest AUROC as shown in Supplementary Fig. 26. In midbrain, the AUROC for our prediction was slightly lower than REPTILE and RFECS, possibly because the DNase experiment performed in midbrain was very noisy; the DNase experiment of mouse e11.5 midbrain was marked as “low SPOT score” in ENCODE, where SPOT stands for Signal Portion of Tag. We found that while 75% to 81% of the genome regions had DNase signals in the other three tissues, only 52% of the genome regions showed DNase signal in the experiment in midbrain. Overall, the comparison shows that our model trained using the *Drosophila* STARR-seq data had similar or better performance than the other methods that were trained directly using mouse experimental data.

For human, we did not have an extensive amount of validated enhancer data. For comparison, we first checked the overlap of our predicted enhancers with the enhancer predictions from chromHMM [55], and SegWay [59], the other unsupervised genome segmentation method, respectively. We observed that a majority of our predictions overlap with either of them (Supplementary Figs. 27-30). In addition, we compared our cell type-specific enhancer predictions with the integrative annotation of ChromHMM and Segway [60] using CAGE-defined enhancers from the FANTOM5 Atlas [61]. The FANTOM5 Atlas has included three human cell lines from the ENCODE project with enhancer predictions from both methods: GM12878, K562 and HepG2. We found that the percentage of overlap for our predicted enhancers was more than three times higher than that of the combined ChromHMM and Segway enhancers in each of these cell lines. Despite the fact that our framework predicted a smaller number of enhancers, the number of overlaps was still higher for our predictions. Around 40% of the CAGE-defined enhancers overlapped with our predicted enhancers, whereas 23% to 34% overlapped with the enhancers predicted by the integrative ENCODE annotation method (Supplementary Fig. 31). We also compared the predicted promoters from our model with their promoter annotations using FANTOM5 promoter sets. Again, the promoters predicted in our model had a higher fraction of overlaps with the FANTOM promoters (Supplementary Fig. 32).

In addition to the integrative ENCODE annotation, we again compared with other supervised enhancer predictions like CSI-ANN [56], DEEP [62] and RFECS [53], using the FANTOM5 enhancer dataset. As most of these methods do not have published enhancer predictions for GM12878 and HepG2, we performed the comparison in the K562 cell line. We found that our predicted K562 enhancers had a similar fraction of overlap with FANTOM5 enhancers compared to that of CSI-ANN, but the fraction was more than twice as high as that of DEEP and RFECS (Supplementary Fig. 33). Besides the overlap with FANTOM5 enhancers, we also checked the overlapping percentage of our predicted enhancers with p300, DHS, and other enhancer binding TFs. Enhancer predictions that have a larger fraction of overlap were regarded as more accurate [53]. For example, RFECS predictions were shown to have a higher positive rate than ChromaGenSVM [63], CSI-ANN [56], and Chromia [53, 64]. Following the same convention, we used their published DNase and ChIP-seq experiments (p300, NANOG, OCT4, and SOX2) in H1-hESCs and repeated the analysis using the same 2.5kb frame distance as they did for overlap. The result showed that our enhancer prediction in H1-hESC had a higher percentage of potentially true positives than RFECS (Supplementary Fig. 34), and thus higher than ChromaGenSVM, CSI-ANN, and Chromia. Overall, our comparisons show that the STARR-seq-based enhancer prediction model is highly accurate in mammalian systems.

### Integrative analysis in human cell lines: Different TFs bind to enhancers and promoters

We further studied the differences in TF binding at promoters and enhancers (Fig. 6 and Supplementary Fig. 35). We focused on the human H1-hESC cell line as there is large amount of functional genomic assays from the ENCODE [52] and Roadmap Epigenomics Mapping Consortium [51] of these cell lines. Together, the consortia have generated ChIP-Seq data for 60 transcription-related factors in the H1-hESC cell line, including a few chromatin remodelers and histone modification enzymes. Collectively, we call these transcription-related factors “TFs” for simplicity.

**Figure 6:**
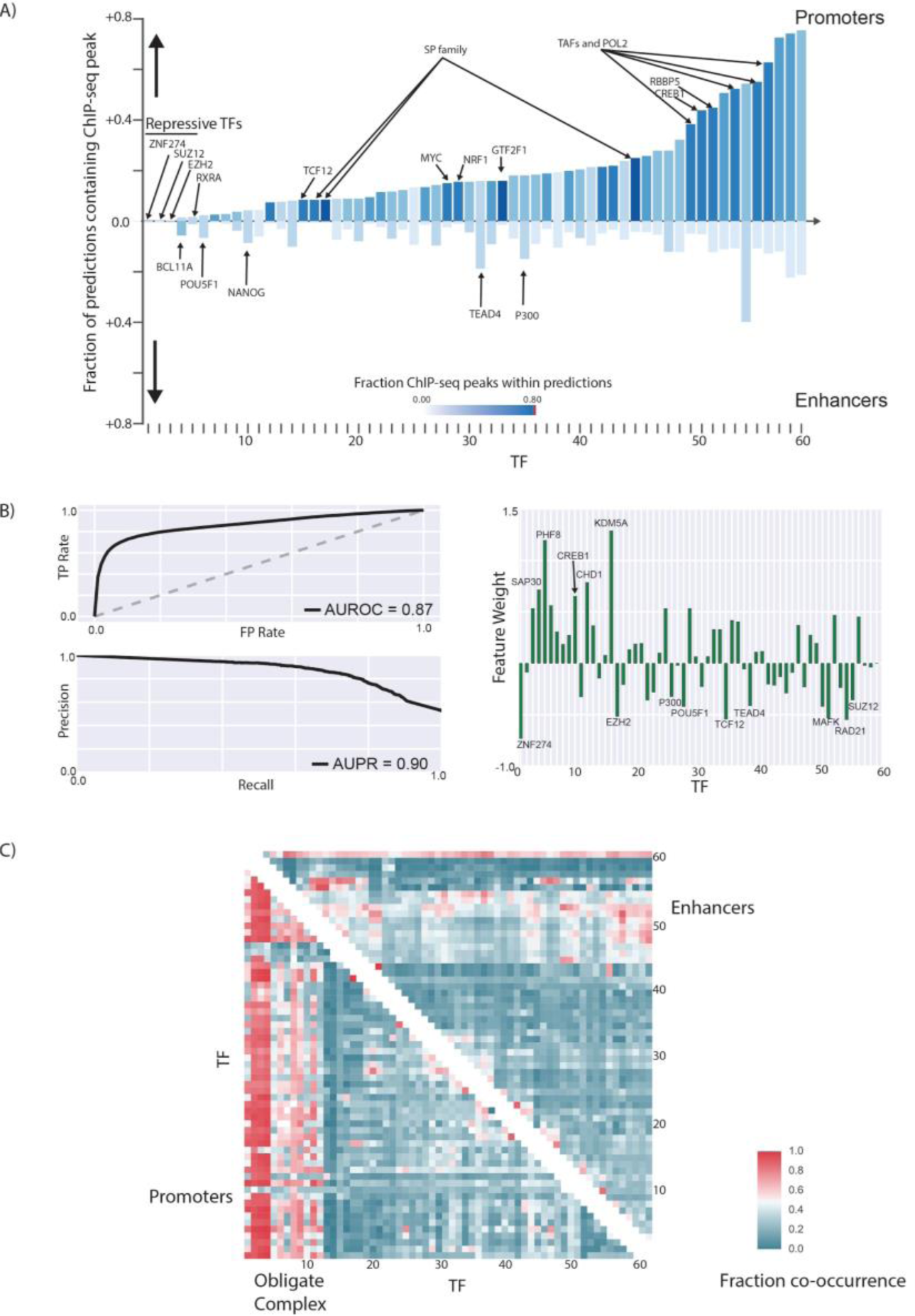
Differences in TF binding patterns at enhancers and promoters. A) The fraction of predicted promoters and enhancers that overlap with ENCODE ChIP-seq peaks for different TFs in H1-hESC are shown. The names of all TFs in the figure can be viewed in Figure S20. B) The AUROC and AUPR for a logistic regression model created using the pattern of TF binding at each regulatory region to distinguish enhancers from promoters are shown. The weight of each feature in the logistic regression model could be used to identify the most important TFs that distinguish enhancers from promoters. C) The patterns of TF co-binding at active promoters and enhancers are shown. The TFs co-occur at promoters regions tend to form obligate complexes. The names of all the TFs in this graph can be viewed in Figure S21.

We showed that the patterns of TFs binding within regulatory regions could be utilized in a logistic regression model to distinguish active enhancers from promoters with high accuracy (AUPR = 0.90, AUROC = 0.87) (Fig. 6). We were also able to identify the most important features that distinguish promoters from enhancers. In addition to TATA box-associated factors such as TAF1, TAF7, and TBP, the RNA polymerase-II binding patterns as well as chromatin remodelers such as KDM5A and PHF8 are some of the most important factors that distinguish promoters from enhancers in H1-hESCs. This provides a framework that can be utilized to identify the most important TFs associated with active enhancers and promoters in each cell type.

We found that although most promoters and enhancers contain multiple TF binding sites, the pattern of TF binding at promoters was different from that at enhancers and that TF binding at enhancers displayed more heterogeneity: more than 70% of the promoters bound to the same set of 2-3 sequence-specific TFs, which was not observed for enhancers (Fig. 6c and Supplementary Fig. 36). The majority of the promoters contained peaks for several TATA-associated factors (TAF1, TAF7, and TBP). These TF co-associations could lead to mechanistic insights of cooperativity between TFs. Similarly, CTCF and ZNF143 may function cooperatively as they are observed to co-occur frequently at distal regulatory regions, consistent with the previous report [65].

To check if the STARR-seq-based enhancer predictions have different TF binding patterns, we referred to the fraction of TF occupancy of predicted enhancer from other methods. The comparison demonstrated in Supplementary Figure 34 shows that the TF binding pattern of our prediction is very similar to that of RFECS. Notably, while RFECS took p300 binding regions as positive training sets, only 25% or less of their predicted enhancers were within 2.5kb of any p300 binding sites, and this is consistent with predictions from ChromaGenSVM [63], CSI-ANN [56], and Chromia [53, 64]. Overall, the high heterogeneity associated with enhancer TF binding is consistent with the absence of a characteristic sequence which can be utilized to identify active enhancers on a genome-wide fashion.

## Discussion

In this study, we developed a framework using transferable supervised machine learning models trained on regulatory regions identified by STARR-seq to accurately predict active enhancers in a cell-type-specific manner. Currently, most existing methods were parameterized on a small number of regions that were typically tested by experiments in an *ad hoc* manner [18, 20, 22-24]. The rich amount of whole-genome STARR-seq experiments established the characteristic pattern flanking active regulatory regions within certain histone modifications [25]. This motivated us to train a shape-matching and filtering model that could be used to identify these patterns in the ChIP-seq signals. As the chromatin marks and epigenetic profiles associated with active regulatory regions are highly conserved among organisms [36-42], we showed that a well-parameterized model in one model organism can be transferred to another with high prediction accuracy.

In the model, we compared close to 30 epigenetic signals for their ability to predict regulatory elements individually. Consistent with previous literature, we found that the H3K27ac matched filter score is very distinctive for predicting active regions, and H3K4me1 and H3K4me3 can distinguish promoters and enhancers. We characterized the amount of redundant information within the metaprofile of different epigenetic features, and showed that the ChIP-seq signals of H2BK5ac, H4ac, and H2A provide independent information that improves the accuracy of promoter and enhancer predictions. In addition to these 30-feature models, we also provide a simple to use six-parameter SVM model for combining H3K27ac, H3K9ac, H3K4me1, H3K4me2, H3K4me3, and DHS to predict active promoters and enhancers in a cell type-specific manner. These six histone marks have been measured for a number of different tissues and cell types by the Roadmap Epigenomics Mapping [51], the ENCODE [52], and the modENCODE Consortia [66]. Using these features, our model could be applied to other species like mouse and human in a tissue- and cell type-specific fashion. We compared our enhancer predictions in mouse and human with the predictions from other published methods. The transferable model trained from the *Drosophila* STARR-seq experiment demonstrated higher performance than the other methods trained directly from mammalian experiments.

While STARR-seq provides a genome-wide unbiased test of the enhancer activity of putative sequences, it is intrinsically episomal and thus not completely revealing the enhancer activity in the native chromatin environment. Selecting for chromosomally active enhancers using H3K27ac and DHS could introduce subtle biases in model training. To address this issue, we employed very different experiment techniques and provided orthogonal validations. This included *in vivo* transgenic assays and *in vitro* transduction assays, in which the predicted regions were tested for regulatory activity in the native chromatin environment. The transgenic assays were performed in e11.5 mice for 133 regions. To better assess our model performance, we also included other published transgenic mouse experiments from the VISTA enhancer browser. In total, we were able to comprehensively validate our tissue-specific predictions in six different tissues in mouse. We also did a similar evaluation with publicly available FIREWACh assay data [30] in mouse, and the result was consistent. With multiple comparisons to other published methods trained directly on mouse data, we showed that the matched filter model is transferable with high accuracy in predicting active enhancers in mouse tissues. The *in vitro* transduction assays were performed in H1-hESCs and three other human cell lines to validate the human regulatory elements predictions. The majority of the predicted elements displayed a significant increase in expression of the reporter gene, further confirming the predictability of our model in mammalian organisms.

As the ENCODE project has generated rich amount of TF ChIP-seq experiment data for H1-hESCs, we used it to look into the differences in the patterns of TF binding at proximal and distal regulatory regions. The TF binding and co-binding patterns at enhancers are much more heterogeneous than that at promoters. This heterogeneity in TF binding patterns makes it more difficult to predict enhancers due to the absence of obvious sequence patterns in distal regulatory regions. However, we were able to create accurate machine learning models that could distinguish proximal promoter regions from distal enhancers based on the patterns of TF ChIP-seq peaks within these regulatory regions. The conservation of the epigenetic underpinnings underlying active regulatory regions sets the stage for our method to study the evolution of tissue-specific enhancers and their genomic properties across different eukaryotic species.

Our results relate to the previous findings that the epigenetic profiles associated with active enhancers and promoters are highly conserved in evolution [36-42]. Therefore, our model of integrating shape-matching epigenetic scores using *Drosophila* STARR-seq enhancers could be applied for prediction in a variety of tissues and cell lines in other species. In the cross-comparison, we showed that the six-parameter integrated model trained with STARR-seq data performs better at predicting mouse tissue enhancers than a model trained in VISTA mouse enhancer data. This highlights the advantage of modeling based on a comprehensive genome-wide experimental assay. In the future, we expect that more extensive whole-genome STARR-seq dataset will become available on mammalian systems. It could be advantageous to re-train the matched filter model on state-of-art datasets. With the setup of our framework, re-training the model with newly generated datasets should be straightforward. We envision that our framework would benefit from these datasets and generate more comprehensive regulatory element annotations across eukaryotic species.

### Implementation: source code and datasets

We have implemented our methods in Python. The source code is available at the website https://goo.gl/E8fLNN. A dockerized image is also provided for download at this site.

The datasets and output annotations referenced in the paper are available in the on the website as well as in the Supplement. The transgenic mouse reporter assay results shown in Supplementary Table 4 and 5 are also made available in the VISTA Enhancer Browser (https://enhancer.lbl.gov). Please refer to the supplement for more details.

## Acknowledgement

M.G. were supported by the NIH grant HG009446-01. A.V. and L.A.P. were supported by NHLBI grant R24HL123879, and NHGRI grants R01HG003988, and U54HG006997, and UM1HG009421 where research was conducted at the E.O. Lawrence Berkeley National Laboratory and performed under Department of Energy Contract DE-AC02-05CH11231, University of California.

